# Pharmacobehavioral space of MoSeq syllables significantly overlaps with scalar locomotion features

**DOI:** 10.64898/2026.08.12.744023

**Authors:** Marti Ritter, Amarender R. Bogadhi

## Abstract

“Revealing the structure of pharmacobehavioral space through motion sequencing” by Wiltschko et al. (2020) has been highly influential in behavioral phenotyping research. In a cohort of nearly 700 mice, the authors demonstrated that Motion Sequencing (MoSeq) could distinguish behavioral effects across a large and diverse set of neuroactive and psychoactive compounds.

A central conclusion of the study is that MoSeq syllable features substantially outperform more traditional scalar behavioral features in treatment classification tasks. Although this comparison is not emphasized outside the Results section, the reported advantage corresponds to an increase in classification performance exceeding 50% relative to scalar feature representations.

While reproducing parts of the analysis using the publicly available dataset, we found that much of this apparent performance difference can be attributed to differences in preprocessing, classifier selection, and hyperparameter optimization. Under alternative, but comparably standard, analytical choices, the performance gap between scalar features and MoSeq syllables was reduced to approximately 11%. Furthermore, in our reanalysis, the performance advantage of MoSeq syllables became statistically significant primarily in highly dense pharmacobehavioral spaces.

These findings do not contradict the utility of MoSeq syllables. Rather, they suggest that the magnitude and generality of their advantage over simpler scalar features may depend strongly on analytical methodology and dataset structure. This distinction is practically relevant, as scalar feature approaches are substantially less computationally demanding and often easier to interpret biologically. Consequently, for laboratories with limited computational resources or for studies focused on specific treatment effects, conventional scalar representations may provide a competitive and more accessible alternative. Our findings highlight the importance of analytical standardization and reproducibility in comparative behavioral representation studies.

## 1 Introduction

Published in September 2020, “Revealing the structure of pharmacobehavioral space through motion sequencing” by Wiltschko et al. describes the ability of MoSeq-extracted syllables to effectively parse the pharmacobehavioral space in a very large dataset of ~700 mice and 16 different treatments (including a control group). As of writing this article, it has 296 citations, and several alternative methods of unsupervised behavior segmentation have been proposed following the original publication (Wiltschko et al. 2015; Luxem et al. 2022). We recently aimed to create a common baseline between the methodology introduced by Datta-lab and those proposed by other groups (Ritter et al. 2026), including VAME (Luxem et al. 2022), SimBA (Goodwin et al. 2024), and A-SOiD (Tillmann et al. 2024). After the observation that none of the compared methods significantly outperformed the others, even including a description limited to aggregated locomotion parameters, we were unsure how to align our results with existing literature.

Because we compared multiple different behavioral summaries with widely varying dimensionalities, we chose a descriptive approach that is as close to the individual model outputs as possible. This included using basic distance metrics (Manhattan-distance) and reducing our application of machine learning to the bare minimum with minimal hyperparameters (i.e. using a Nearest Centroid classifier) to avoid the inadvertent introduction of bias of model selection. This is a deviation when compared to the analysis applied by Wiltschko et al., who utilized cosine distances to describe the topology of feature spaces and applied a set of linear classifiers (Logistic Regression & Support Vector). The logistic regression classifiers were optimized using a regularization parameter chosen from within a search space to obtain the best results for a scalar and a MoSeq-derived feature set. Further hyperparameters were also shared.

We initially aimed to reproduce a set of distance-based descriptions included in Figure 4 of Wiltschko et al. (2020). A ratio of distances (Cosine or Manhattan) was chosen as a first comparison as this is a metric common between both studies. As we attempted to reproduce their outcome, we realized that the distance ratio featured in Fig. 4B did not match our expectations based on the other metrics shown in the same panel. While between-cluster distances were greater for MoSeq-derived features, so were within-cluster distances. Wiltschko et al. also acknowledge this and interpret the increased distances as behavioral variance between individuals.

**Figure 1.**
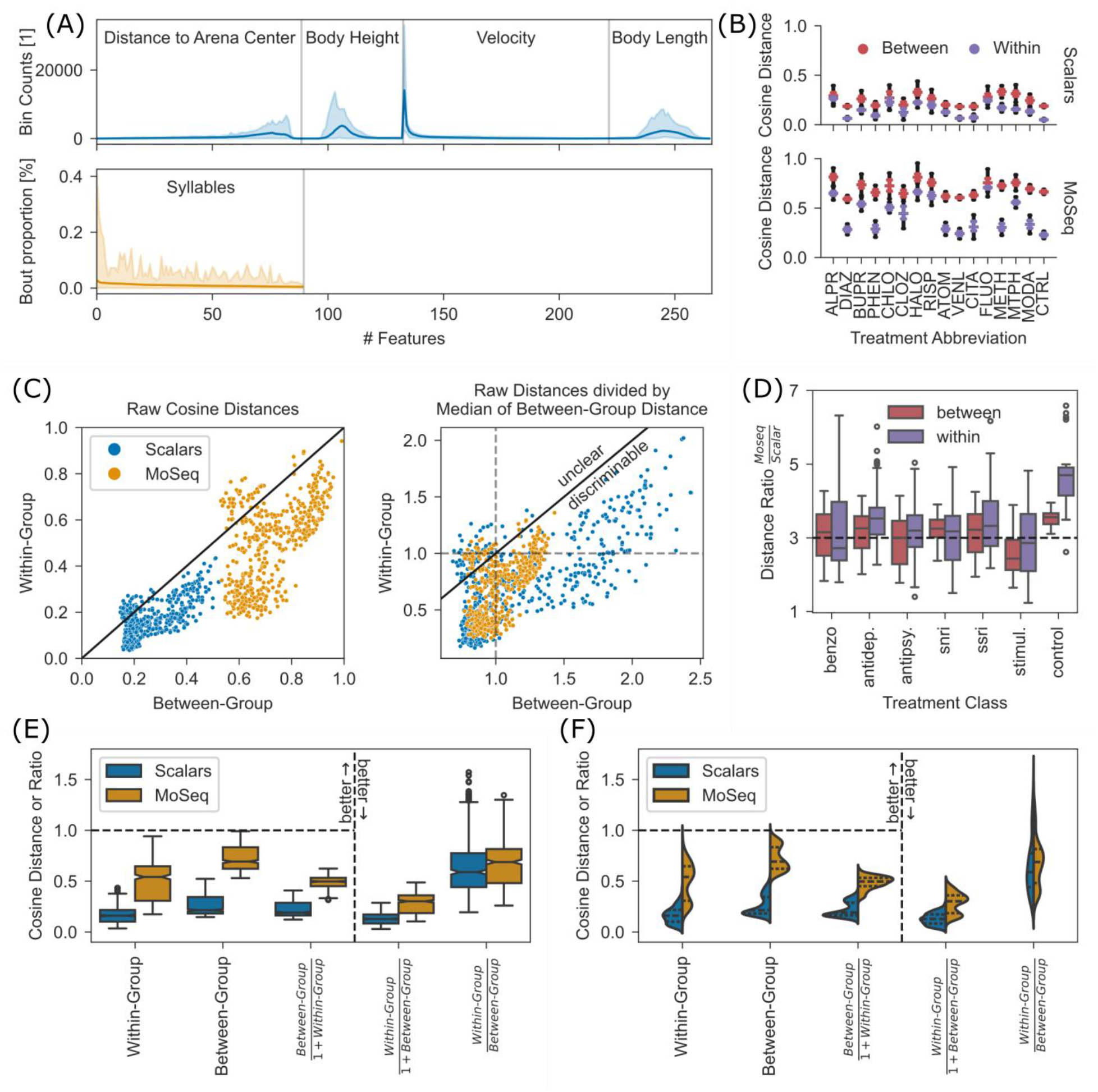
Overview of feature set topology. (A) Comparison between distributions of scalar (top) and MoSeq-derived (bottom) features. Scalar features were computed as a concatenation 4 locomotion parameter histograms, MoSeq-derived features were calculated as the bout proportion of 90 different behavioral syllables. (B) Reproduction of Wiltschko et al.’s Figure 4a, based on released data. (C) Scatterplot showing relationship of average cosine distances between samples of different groups (“Between-Group”) and samples of the same group (“Within-Group”). Left plot shows raw distances; right plot corrects for between-group distance by dividing both axes by its median. (D) Boxplot of the ratios between distances found in scalar and MoSeq-derived feature sets, grouped by drug class and distance type (between or within group). (E) Extended reproduction of Wiltschko et al’s Figure 4b. Bottom left box shows ratios present in Wiltschko et al. Boxplots right of the dashed line are additional measures. Plots left of the line have their optimal value at 1 (optimum-1), while plots to the right show optimum-0 metrics. All comparisons are significant (two-sided t-test, Bonferroni correction, p<=0.05). (F) Violinplot of the same data as in panel E. None of the presented distributions follow normality.

**Figure 2.**
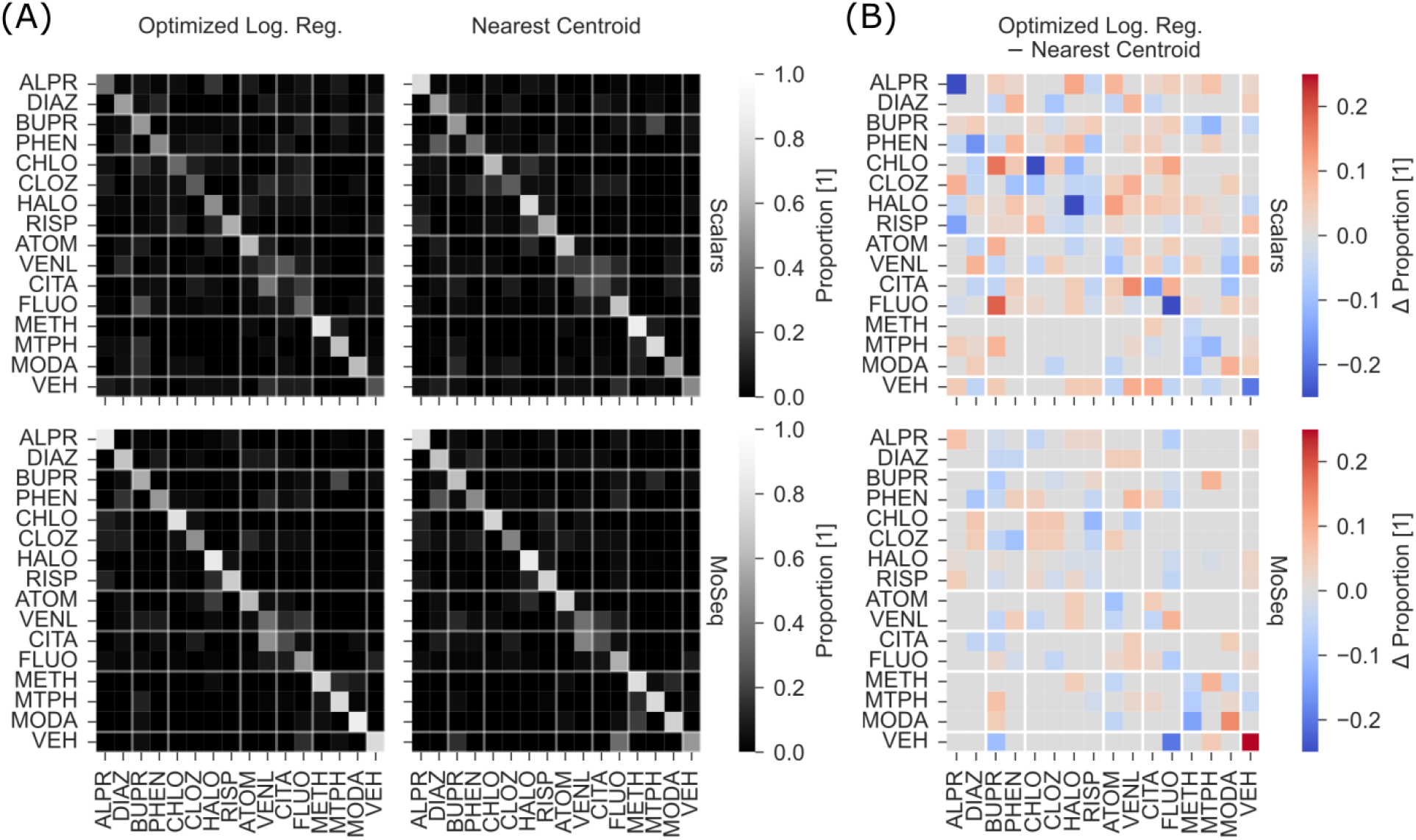
Confusion matrices for two classifiers. (A) Left column shows the inter-treatment confusion matrix for two optimized Logistic Regression classifiers applied to both feature sets; right column shows the same confusion matrix for Nearest Centroid classifiers. (B) Difference between the left and right column of A for both feature sets.

**Figure 3.**
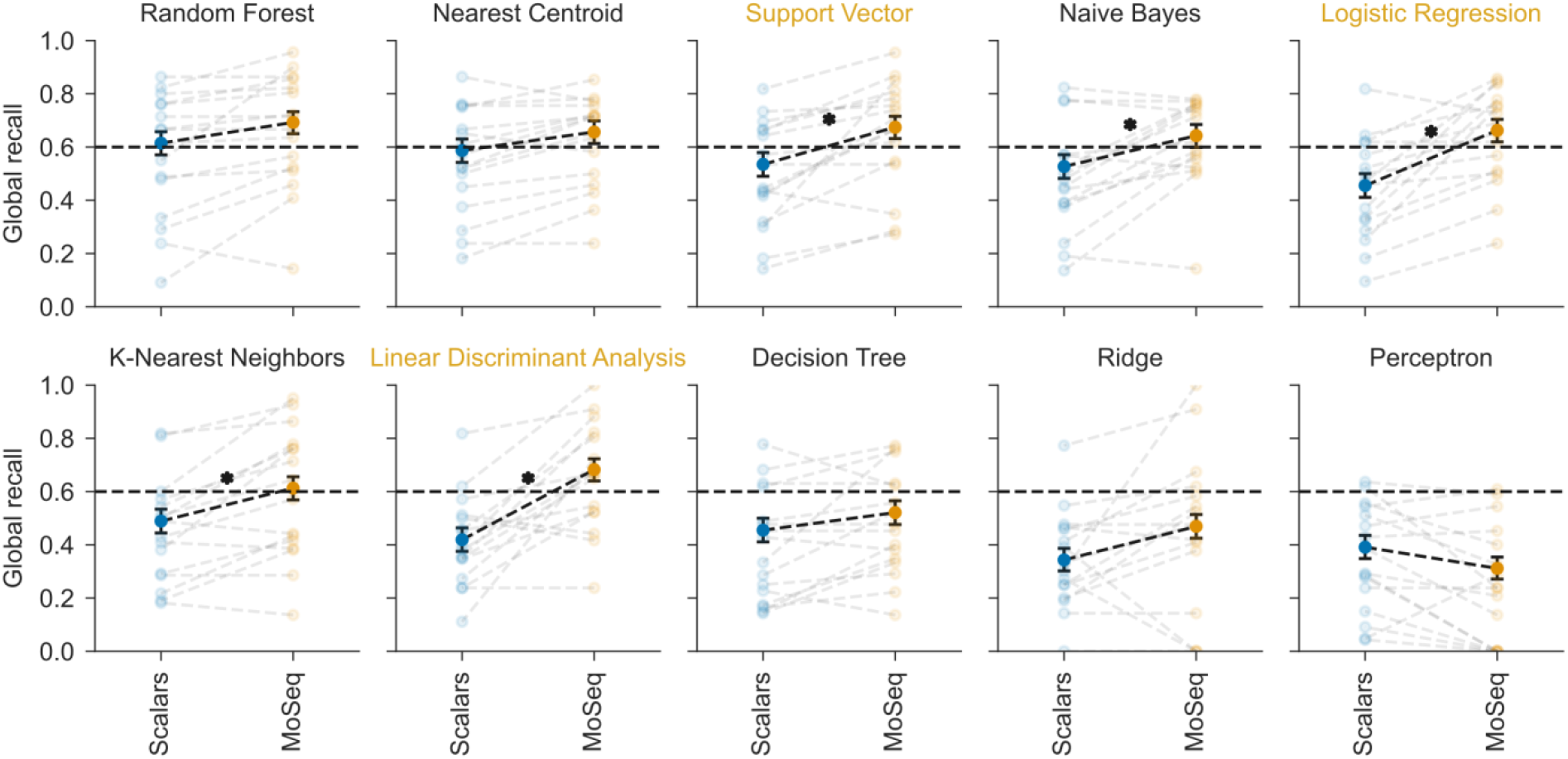
Global recall and binomial statistics for both feature sets across multiple classifiers. Ten classifiers were applied to both feature sets. Point plots of global recall (CI=95%) are shown, along with treatment-wise statistics shown in background. Significant differences between feature sets are marked with an asterisk (two-sided WSR, Bonferroni correction, p<=0.05). Order of classifiers was chosen based on the average global recall across scalar and MoSeq-derived feature sets. Classifiers marked with color are featured in Wiltschko et al.

**Figure 4.**
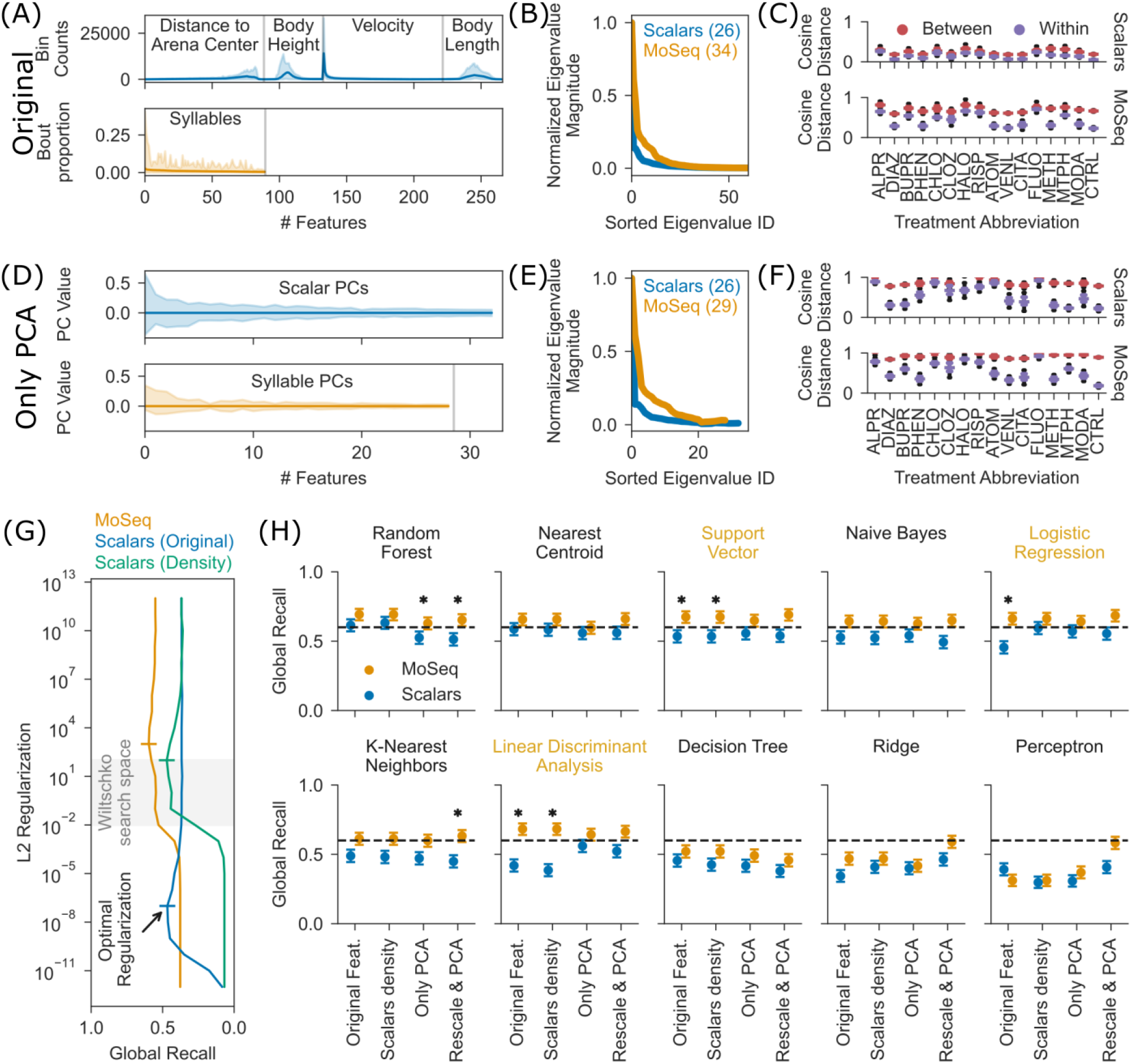
Effects of preprocessing on topology and classifier performance. (A) Reproduction of Figure 1A to allow for easier comparison. (B) Reproduction of Wiltschko et al.’s Extended Data Fig. 4c. (C) Reproduction of Figure 1B to allow for easier comparison. (D) Feature distribution after application of PCA on features shown in panel A. Scalar features were rescaled to density rather than counts before PCA was applied. (E) Same layout as panel B but applied to PCA-transformed features. (F) Same layout as panel C but applied to PCA-transformed features. (G) Effects of L2 Regularization on global recall in a logistic regression classifier, based on three different feature sets: MoSeq syllable proportions, raw scalar histogram counts, and scalar histogram density distribution. Optimal regularization parameters (maximum global recall) shown as horizontal marker, and search space featured in Wiltschko et al. shown as horizontal gray span. (H) Effects of preprocessing on classifier performance. Same layout as in Figure 3, but split across possible preprocessing steps. From left to right: Raw features as shown in Figure 3, scalar histograms transformed to densities rather than counts, PCA applied to syllable proportions and scalar densities without prior standard scaling, and finally PCA with preceding standard scaling applied feature-wise. Significant differences in global recall between scalar and MoSeq-derived features are marked with an asterisk (two-sided WSR test, paired on treatment recall, Bonferroni correction, p≤0.05).

Yet, further analysis of the data shared publicly by Wiltschko et al. shows that the underlying topology of both feature sets is very similar, and that the primary difference in cosine distance between them is caused by scaling. Consequently, the significant differences in performance between scalar- and MoSeq-derived features across classifiers cannot depend solely on a significant increase in meaningful information in the MoSeq syllables over scalar aggregation of locomotion parameters. These additional analyses and findings, based on data publicly shared by Wiltschko et al., reveal significant limitations to the interpretation found in their publication.

## 2 Materials and Methods

### 2.1 Data sources

In short, Wiltschko et al., applied a large set of treatments (15 treatments of various drug classes + 1 control group) to between 6 and 12 mice per group for a total of 673 animals according to their methods. All data described in this chapter relates to 6–8-week-old C57BL/6 males supplied by Jackson Laboratories.

All mice were recorded in a circular arena that was filmed from the top by a camera capable of recording depth maps. In contrast to our previous study, Wiltschko et al. were thus able to apply MoSeq rather than Keypoint-MoSeq. To test their primary hypothesis of an improved separation of treatment groups in a MoSeq-derived feature space, they calculated the histogram of a set of locomotion parameters and concatenated the resulting bin counts. Two different linear classifiers (Logistic Regression and Support Vector) were fit on each of these feature sets and their performance (measured by an F1-score) compared. A later visualization (Figure 5d) also applied a Linear Discriminant Analysis. For further details on their methods, please see Wiltschko et al. (2020).

**Figure 5.**
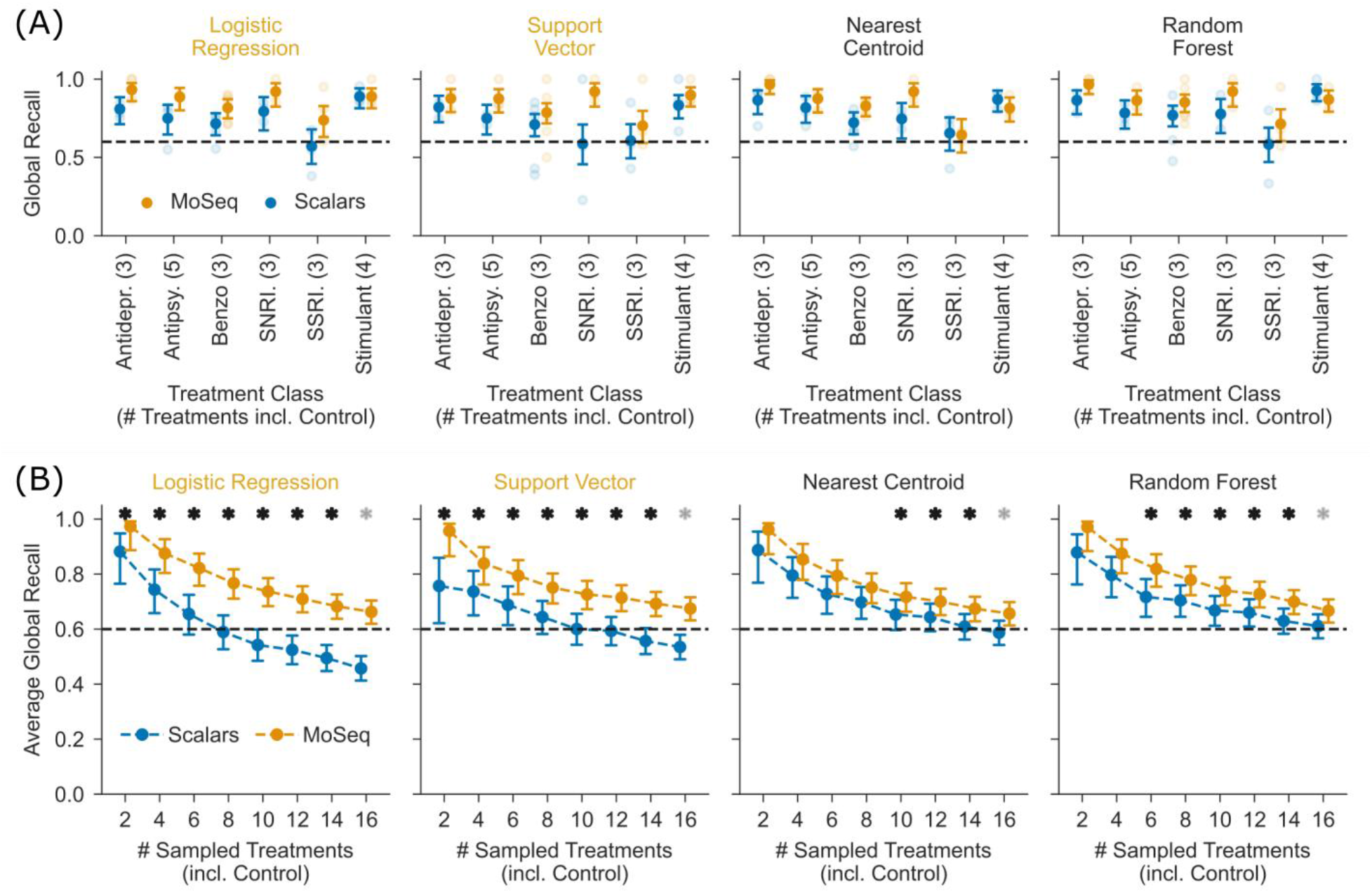
Effects of pharmacobehavioral space density on classifier performance. (A) Point plots of global recall with same properties as Figure 3, but with subset-wise statistics removed for readability. A selection of four classifiers (best performing two from Figure 3 and two featured in Wiltschko et al.) were applied to feature sets limited to 1 of the 6 treatments classes featured in Wiltschko et al.. Each one included the control group. None of the comparisons between feature sets were significant (two-sided WSR test, paired on treatment recall, Bonferroni correction, p≤0.05), due to the limitation to only a small number of treatment groups. (B) Extension of panel A to randomly sampled treatment subsets of increasing length (see Results). Again same layout as Figure 3. All subsets included the control group. Significant differences in average global recall are marked with an asterisk (two-sided MWU test, Bonferroni correction, p<=0.05). Transparent asterisk for n=16 represents an estimate for single possible combination of all treatment groups, assuming significance when n=14 was already significant.

As only 501 of the mice can be found in the shared data (doi.org/10.5281/zenodo.3951698, version 1.0, file “fingerprints.pkl”) and their Supplementary Table 1, we are uncertain what experiments the remaining 172 animals were used in. If we assume that the 95 animals of a different strain described in a later analysis were included in this number, this still leaves 77 mice we cannot assign to a particular experiment. Regardless of this question, we analyzed the available dataset of 501 mice and their respective scalar and MoSeq-derived features.

### 2.2 Statistical Analysis

All statistical analysis was performed in Python 3.11.9. Unless stated otherwise we used a Wilcoxon signed-rank (WSR) test, and significance levels set at α=0.05 with a Bonferroni-correction based on the number of tests visible in each plot.

#### 2.2.1 Two-way ANOVA

The `ols` class and `stats.anova_lm` function implemented in the statsmodels library (version 0.14.2) was used to apply a two-way type 2 ANOVA to the treatment recall shown as background plots in Figure 3. The applied formula was `“treatment_recall` ~ `C(classifier) * C(feature_set)”`.

#### 2.2.2 Cosine distance

The cosine distance corresponds to the inverse of the cosine similarity, and as such is a measure of angular distance between two points in multidimensional space. In this chapter we use cosine distances, rather than Manhattan distances, to stay in line with previous analysis done on this dataset.

The equation for cosine distances is 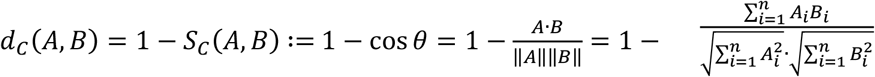, with A and B being two points in n-dimensional space, and *S*_*C*_ representing cosine similarity.

#### 2.2.3 Paired T-test

We used SciPy’s (version 1.14.0) implementation of the paired t-test (function `ttest_rel`) in the following analysis:

- In Figure 1E&F a two-sided test was used to stay in line with the analysis shown in Wiltschko et al. All comparisons were significant at the given significance level and correction.

#### 2.2.4 Shapiro-Wilk test

We used SciPy’s (version 1.14.0) implementation of the Shapiro-Wilk test for normality (function `shapiro`) in the following analysis:

- In Figure 1E&F we were able to show that none of the observed distributions followed normality at the given significance level and correction.

#### 2.2.5 Wilcoxon signed-rank test

We used SciPy’s (version 1.14.0) implementation of the Wilcoxon signed-rank (WSR) test (function `wilcoxon`) in the following analysis:

- To account for the lack of normality of the distributions shown in Figure 1E&F, we followed up the previous two-sided paired t-test with a two-sided WSR test. All comparisons remained significant.
- In Figure 3 we applied a two-sided test to test significant differences in performance between scalar and MoSeq-derived features within the same classifier.
- To evaluate the possible effect of feature preprocessing on classifier performance we applied a two-sided version of this test in Figure 4H.
- In Figure 5A we applied a two-sided test to verify if classifiers performed significantly differently on scalar and MoSeq-derived feature sets when only applied to single treatment classes, rather than the full set of treatments. None of the tested comparisons were significant at the given significance level and correction.

#### 2.2.6 Mann-Whitney U-Test

We used SciPy’s (version 1.14.0) implementation of the Mann-Whitney U-Test (function `mannwhitneyu`) in the following analysis:

- In Figure 5B we tested the performance of classifiers on significant differences between scalar and MoSeq-derived features across randomly sampled combinations/subsets of treatment groups with sizes between 2 and 16 (always including the control group). Due to the possible repeated sampling of the same treatment group in two separate subsets of same size, we decided to avoid a paired test and chose the MWU in place of the usual WSR test.

#### 2.2.7 Bonferroni correction

All statistical tests that were applied to more than one comparison were corrected for false-positive discovery by applying the Bonferroni correction. Unless stated otherwise we used the significance threshold of α=0.05 and corrected according to **α**_**Bonferroni**_ **= α**_**uncorrected**_**⁄n**_**tests**_, with **n**_**tests**_ corresponding to the number of tests performed (nearly always the number of tests shown in a plot).

We applied this correction to the paired t-test and subsequent Shapiro-Wilk and WSR test in Figure 1E&F (**n**_**tests**_ = 5); the WSR test in Figure 3 (**n**_**tests**_ = 10); the WSR test in Figure 4H (**n**_**tests**_ = 40); the WSR test in Figure 5A (**n**_**tests**_ = 24); and the MWU test in Figure 5B (**n**_**tests**_ = 32).

#### 2.2.8 Binomial confidence intervals

To remain in line with the visualizations shown in the previous chapter, global recall of the tested classifiers/feature set was marked with error bars showing the confidence interval of the underlying binomial result: Samples were either assigned correctly or not. We calculated the test statistic (recall) using SciPy’s binomial test function (function `binomtest`) and translated it to a confidence interval using the normal distribution implementation (function `norm`). The test statistic and interval are shown in Figure 3, Figure 4H, and Figure 5A&B. In contrast to our previous analysis, we did not evaluate performance above a particular chance level but compared performance between the two feature sets.

#### 2.2.9 Intrinsic dimensionality by eigenvalue magnitude

Wiltschko et al. use the algorithm proposed by Fukunaga and Olsen (1971) to estimate the effective or intrinsic dimensionality of both feature sets. As described by them this includes the eigenvalue decomposition of the covariance matrix of a given feature set, followed by the normalization of eigenvalue magnitudes to the range between 0 and 1. The number of rescaled magnitudes above a certain cutoff, given as 0.01, represents the effective dimensionality of the source feature set.

### 2.3 Classification

To account for the effect of classifier selection, feature preprocessing, and hyperparameter selection, we used mostly the Scikit-Learn Python package, version 1.5.1.

#### 2.3.1 Feature pre-processing

Data shown in Figure 1, Figure 2, Figure 3, and Figure 5 was not further processed and includes only those features shared by Wiltschko et al. publicly. These features are four concatenated count histograms that make up the scalar feature vector, and the bout proportions of 90 syllables in the MoSeq feature vector. In contrast to their claim in the methods, Wiltschko et al. did not bin each feature (Position, Height, Velocity, Length) into 90 bins each, but applied 90 borders, so that the resulting 89 bins per scalar property lead to a vector of length 266, rather than the expected 270 features.

In Figure 4 we applied preprocessing steps to verify whether the performance of classifiers trained on both features was affected. As a preceding step we transformed the bin count histograms to density histograms using NumPy (version 1.26.4, function `histogram`). The two following preprocessing steps, performing a PCA on raw features and performing a PCA with standard-scaled features, were performed using Scikit-Learn and its classes `PCA` and `StandardScaler`.

#### 2.3.2 Classifiers

From Figure 1 onwards we applied a selection of classifiers available from Scikit-Learn. `LogisticRegression`, `SVC` (Support Vector Classifier), and `LinearDiscriminantAnalysis` are also featured in Wiltschko et al, and we extended this selection by `RandomForestClassifier`, `NearestCentroid` (based on Manhattan distance), `GaussianNB` (Gaussian Naïve Bayes), `KNeighborsClassifier`, `DecisionTreeClassifier`, `Ridge`, and `Perceptron`. Logistic Regression, Support Vector, and Perceptron classifiers are innately binary classifiers and were consequently translated into multi-class classifiers using the `OneVsRestClassifier` adapter. Where applicable we set a fixed random seed of 0 to ensure reproducibility, and used the same parameters as described by Wiltschko et al.. Every other parameter was left at the defaults present in version 1.5.1 of the Scikit-Learn Python package.

#### 2.3.3 Cross-Validation

In contrast to the `StratifiedShuffleSplit` applied by Wiltschko et al. (500 times sampled, 10% of data held out per sample), we applied a `LeaveOneOut` cross-validation strategy. With 501 mice in the available dataset this leads to 501 possible test-sets, corresponding to a 501-fold cross-validation. With each fold one prediction label per test sample is returned, and all following analysis (recall, confusion matrix, etc.) are then run on the combined cross-validated prediction labels. While Wiltschko et al. refers to their strategy as a 500-fold cross-validation, the name available in Scikit-learn is more appropriate, as a fold represents a subset of data and each sample is conventionally present in one and only one fold.

#### 2.3.4 Hyperparameter search

Logistic regression classifiers can apply an L2 weight penalty based on an inverse regularization strength C. Wiltschko et al. scanned a logarithmic range of 5 values for C from 0.01 to 100 for both feature types and presented the optimal choice per feature. As we observed no visible improvement in performance for scalar features across that range, we extended the search space from 10^−12^ to 10^12^ for both MoSeq and scalar features, as well as the density-transformed scalar features, and presented the outcome of that choice in Figure 4G&H.

## 3 Results

After we found the distance ratios presented by Wiltschko et al. to deviate from our expectations, we retraced parts of their analysis in search of common ground between their results and the results of our most recent research (Ritter et al. 2026). We downloaded the tracking and feature data that they shared on Zenodo (see Methods) and applied similar methods to theirs, adding analysis that could bridge the gap between their primary findings and ours. For basic details on their methodology and data acquisition see Methods (section Data sources), or for more detail see Wiltschko et al. (2020).

### 3.1 Cosine distances in scalar and MoSeq feature sets have different scales but similar topology

Scalar locomotion-derived histograms and the MoSeq-derived syllable proportions are very distinct feature sets, as can be seen in their length and distribution of variance (see Figure 1A). While their respective cosine distances appear at first dissimilar (see Figure 1B), a comparison of sample-wise averages of within- and between-cluster distance spaces shows in fact a similar topology (see Figure 1C). Cosine distances in the MoSeq-derived feature set are usually around 3 times larger than those found in the scalar feature set (see Figure 1D). Because of this core similarity, the pure ratio between within and between distances remains very similar, as follows from 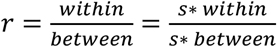. But while the ratio present in Wiltschko et al. is given in their methods by 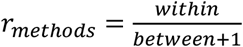, this does not match the results shown in their Figure 4B, based on measuring the averages shown in the boxplots. Average distances shown for the scalar feature set are roughly 0.084/0.225 (within/between), suggesting an r≈0.069 but shown as roughly 0.208. Similarly, for syllable features distances shown are roughly 0.287/0.69, suggesting an r≈0.17 but shown as roughly 0.536. The ratio boxplots can be reproduced by applying 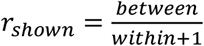 instead (see Figure 1E). The most significant effect of this change is the inversion of the “optimal” ratio from 0, due to 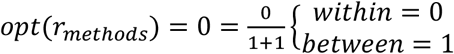, to 1 based on 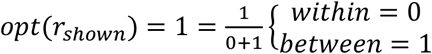. Qualitatively both ratios still do not differ much, as the inclusion of an additive 1 in the denominator significantly affects the outcome, especially for smaller values. Independent of the choice of ratio, all distances and ratios are significantly greater for MoSeq-derived features when compared to scalar-derived features (two-sided t-test, Bonferroni correction, p<=0.05). While the choice of a t-test in this evaluation is not recommended due to the lack of normality in the distance and ratio distributions (Shapiro-Wilk test, Bonferroni correction, p<=0.05) we still applied it to remain in line with previous analysis (see Figure 1F). Replacing the t-test with a Wilcoxon signed-rank (WSR) test did not change the outcome (two-sided WSR test, Bonferroni correction, p<=0.05).

These findings suggest that the underlying topology of both feature sets is very similar, and that the primary difference in cosine distance between them is caused by scaling. This in turn would not affect the relative behavior of distances within each feature set but does so in the ratio chosen by Wiltschko et al. due to the choice of a large additive to the denominator, to the degree that inverting enumerator and denominator has only a minor qualitative effect on the ratio.

### 3.2 Choice of classifier significantly affects discriminative performance on scalar feature set

As the topology within each feature set appeared similar, we were interested to see whether the choice of classifier led to the difference in findings between Wiltschko et al. and our most recent study (Ritter et al. 2026). The linear Logistic Regression classifier chosen in Wiltschko et al. presents a significant improvement in performance in the MoSeq feature set over the scalar feature set (compare left column of Figure 2A). Yet, when we chose a Nearest Centroid classifier, using the same 501-fold cross-validation for both classifiers (see Methods), there seemed to be no significant difference across behavioral summary models (compare right column of Figure 2A). Our choice of this classifier was motivated by the wish to avoid choosing hyperparameters, and to remain as close as possible to the underlying topology of the features produced by each summary model. The sole choice in a Nearest Centroid is the selection of the distance metric. In our case we used Manhattan distance, as it has been shown to be less affected by increasing dimensionality (Aggarwal et al. 2001) and evaluates features of different type and scale equally.

It is apparent that Logistic Regression performs worse on scalar-derived features than the Nearest Centroid classifier (compare top row of Figure 2A), with both having a similar recall on MoSeq-derived features (compare bottom row of Figure 2A). Visualizing the specific changes in classification proportions shows that most of the different outcomes between both classifiers are misassignments of the Logistic Regression classifier in the scalar feature set, visible as reduced assignment proportion along the diagonal of the confusion matrix (see Figure 2B). Curiously, this reduction in performance seems focused on few treatment groups (Alprazolam, Chlorpromazine, Haloperidol, and Fluoxetine) across several treatment classes. In the MoSeq feature set, the logistic regression classifier performs better when separating control group animals from Fluoxetine.

This outcome suggests that there exists a way to meaningfully discriminate treatment groups even within a scalar feature space, as otherwise it would be not possible to successfully apply any classifier to this task.

### 3.3 Significant difference in performance Scalar vs MoSeq depends on classifier choice

Since the choice of the classifier clearly matters, we arbitrarily selected 10 of the classifiers available from Scikit-Learn and fitted all to both feature sets. All classifiers were cross-validated using the same 501-fold approach (Leave-One-Out, see Methods). Sorting the resulting classifications by average global recall across samples and feature sets shows that the best performing classifiers (Random Forest and Nearest Centroid) are able to predict on both feature sets with only insignificantly worse global recall when trained on scalars features (see Figure 3). The classifiers and methods featured in Wiltschko et al. can be found mostly in the mid-range of global recall (0.55-0.60) and all mid-range classifiers present significantly worse recall when trained on scalar features. It also appears that classifiers in the lower recall range cannot work well on the present features, independent of the feature set.

A two-way ANOVA further confirms these results and reveals a significant main effect for both classifier choice (F(9, 300)=7.56, p<1e-9) and choice of feature set (F(1, 300)=22.77, p<1e-5) on treatment recall, with no significant interaction between classifier and feature set choice (see Methods).

This finding suggests that the significant difference in performance for scalar- and MoSeq-derived features across classifiers cannot depend solely on a significant increase in meaningful information in the MoSeq syllables over scalar aggregation of locomotion parameters.

### 3.4 Cosine distances and classifier performance significantly depend on feature preprocessing

Since the difference in performance for MoSeq-derived and scalar feature sets across the selection of classifiers cannot be explained solely by information available in only the MoSeq-derived features, we investigated the effects of common preprocessing steps on global recall. It is well known that optimization-based linear classifiers (i.e. Logistic Regression, Support Vector, LDA) are sensitive to feature scaling and sometimes require preprocessing of features. This can be argued to originate in the objective of modelling class boundaries with a single function, rather than single points (Nearest Centroids) or multiple independent separation layers (Random Forest).

Wiltschko et al. argue that the effective dimensionality of MoSeq-derived features, and as such its intrinsic capacity to describe behavioral variability, is higher than that of scalar-derived features. We can recreate their findings on the shared dataset (see Figure 4A&C), resulting in the same number of intrinsic dimensions by using the algorithm proposed by Fukunaga and Olsen (1971) as shown in Wiltschko et al. (see Figure 4B).

Prior to the application of a PCA, scalar features were transformed into densities, rather than bin counts, as we wanted to avoid further feature-wise rescaling at this point to not lose the inter-feature relationship present in the histograms. Applying a PCA limited to the number of components needed to explain 95% of variance in each feature set results in a significant reduction of features (see Figure 4D). While this transformation should in theory maintain the majority of variance in each feature set, it still results in a reduction of intrinsic dimensionality according to the Fukunaga-Olsen estimate used by Wiltschko et al. (see Figure 4E), and reduces the intrinsic dimensionality gap between scalar and MoSeq from 8 to 3.

Interestingly, this transformation also visibly narrows the difference in intra- and inter-group cosine distances between scalar and MoSeq feature sets (see Figure 4F). This appears not to be caused by a reduction of cosine distances within the MoSeq feature set, but rather an increase of cosine distances in the scalar features.

While preparing the scalar features for the PCA by rescaling them to densities, we noted that the performance of the Logistic Regression classifier significantly improved. This appears to have been related to the search space used in Wiltschko et al. (see Figure 4G). While they performed the search for the L2 regularization parameter within a logarithmic range between 10^-2^ and 10^2^, a search between 10^-12^ and 10^12^ yields not only the optimum for the scalar feature set (found at 10^-7^), but also the optimum for the MoSeq-derived feature set (set to 10^2^, is found at 10^3^). Translating the scalar features to densities rather than bin counts leads to their optimal regularization being found at 10^2^.

To finalize the usual preprocessing pipeline found in many machine learning applications, we also applied a feature-wise standard-scaling (mean to zero, standard deviation to one; transformation to z-score) followed by again a PCA (see Methods). Across all four preprocessing stages (no preprocessing, scalars to densities, PCA without rescaling, and rescaling followed by PCA) some classifiers improved their global recall, and some became worse, but across all three linear classifiers featured in Wiltschko et al. there was a point beyond which the difference in performance on scalar and MoSeq-derived features stopped being significant (see Figure 4H).

These findings underscore the sensitivity to feature scales of linear classifiers, and the need to preprocess datasets. Furthermore, the finding of unchanged inter- and intra-cluster cosine distances in MoSeq features combined with a reduction of effective dimensionality after a PCA (without rescaling), suggests that the PCA for 95% variance filtered out dimensions that captured individual rather than treatment-related variance. Since the usual goal of this analysis is to find treatment-specific rather than individual changes in behavior, this loss in resolution could improve interpretability of the results.

### 3.5 Differences in performance only become significant in dense pharmacobehavioral spaces

While the significant differences in performance between scalar and MoSeq-derived feature sets disappear after some basic preprocessing of the original data, it was clear that the dataset collected by Wiltschko et al. contained a very high number of treatment groups and classes compared to our previous study (Ritter et al. 2026; 9 treatment-dosage groups, including 1 control). It might be that we were unable to see differences in performance across our behavioral summary models due to the limited number of treatments and mice: 3 treatments, 2 of which were stimulants and 1 potential treatment for major depressive disorder, with 31-48 animals per group and control.

Separating Wiltschko et al.’s unprocessed dataset by treatment class showed that the global recall of four selected classifiers (2 featured in Wiltschko et al. and the two best from Figure 3) performed far better on small sets of treatment groups when compared to the full set (see Figure 5A). Additionally, the significant differences of linear classifier models between scalar and MoSeq-derived features were not apparent within the subsets.

Extending this analysis to its logical conclusion, we sampled up to 15 combinations of various lengths (1-15) from the 15 available treatment groups and extended each with the control group (see Figure 5B). For example, a length of 3 allowed for 455 possible combinations and we sampled 15 of these. The 15 possible combinations of length 1 on the other hand corresponded to the 15 treatment groups (one group per combination), and length 15 allows a single possible combination containing all groups (see legend Figure 5B regarding transparent asterisk). This analysis reveals that while the difference of global recall averaged across all sampled combinations between scalar and MoSeq-derived features is already significant for both linear classifiers at a single treatment and control group, it only becomes significant in pharmacobehavioral spaces with more than 6 to 10 treatment and control groups for Random Forest or Nearest Centroid classifiers. This appears to not be generally caused by a faster decay of performance on the scalar feature set, but rather the decrease in variance of global recall as more treatment groups are sampled.

These results suggest that the choice of a classifier is central to the evaluation of scalar or MoSeq-derived features. In addition, there appears to be a comparatively large range of possible treatment group configurations in which the difference in recall between scalar and MoSeq features remains non-significant. Thus, other factors, such as the explainability of effects or reproducibility of features, should take precedence over raw classification performance for low-density pharmacobehavioral spaces or certain treatment classes.

## 4 Discussion and conclusion

The introduction of data-driven behavioral segmentation methods opened the door to a deeper, more nuanced analysis of the rich phenotypes present in both natural behavior (Ritter et al. 2025) and the effects of interventions or disorders (Gschwind et al. 2023). Possible approaches include the analysis of recurrent patterns (i.e. VAME, Luxem et al. 2022) or the frequency of transitions between them (behavioral flow analysis, Ziegler et al. 2023). One of the most known publications in this field is “Revealing the structure of pharmacobehavioral space through motion sequencing” by Wiltschko et al. (2020), cited at time of writing around 300 times (compared VAME’s roughly 180 citations and behavioral flow analysis’ 19). This publication also had a strong influence on our study planning and our previous goal of finding the underlying syllable structure of social behavior in mice. Consequently, we were surprised when the results of our second study (Ritter et al. 2026) did not match expectations, i.e. lack of significantly increased discriminative performance of unsupervised behavioral summary methods over mere aggregation of locomotion parameters.

As we investigated the cause of the divergent results, we unexpectedly discovered multiple points separating our analysis from the one shown by Wiltschko et al. These points covered choices of measures (see Figure 1E,F), potential decisions regarding preprocessing (see Figure 4A-F, H), and classifier and hyperparameter choices (see Figure 2, Figure 3, and Figure 4G). Cumulatively these aspects had a significant effect on the outcome of both analyses. For example, judging by Figure 3D in Wiltschko et al. Scalars performed with a global F1-metric of around 0.4, while MoSeq reached F1≈0.62. This gap suggests a nearly 55% improvement in F1 when choosing MoSeq over scalar features. We can reproduce this gap, with scalar features allowing an F1≈0.45 and MoSeq features offering F1≈0.66. Yet, when applying a nearest centroid classifier on the same data, this increases to F1≈0.59 for scalar features, while MoSeq’s performance remains unchanged. This reduces the estimated performance gain of choosing MoSeq over scalar features to around 11%. Additionally, it appears that the impressively high number of treatment groups found in their foundational study could be required for the choice of MoSeq-derived syllables over scalar features (see Figure 5). Based on the deep methodology and high investment of time and animals found in their study, we expected that these alternative avenues were exhaustively analyzed.

Our reproduction of their results is potentially limited, as we could not find data for between 77 to 172 animals mentioned in their methods, and cannot know whether the shared data was further processed beyond what we understood from the published text. Consequently, it is possible that the quantitative deviations in our output are due to our limitation to available data. Nevertheless, we followed the publication as closely as possible regarding parameter choices and the reproduced figures appear to qualitatively align with the published results, so that the conclusions drawn here are most likely valid.

Still, parts of the results shown in the publication are not accessible to us, as we do not have access to the required data. This includes the analysis of on- and off-target effects in syllable expression in treated mouse models of autism spectrum disorder. It would interest us to see the same output reproduced with scalar features or their corresponding principal components. Theoretically, it would be just as valid to select scalar locomotion features rather than syllables as treatment targets and could even offer improved explainability.

Another important difference is found in the applied cross-validation strategies. Wiltschko et al. selected a stratified and shuffled 500-times split of their data, with each shuffled split containing 10% held-out data and 90% training data. As we had already chosen a leave-one-out (here 501-fold cross-validation) approach in our previous study, we chose to maintain that decision. Qualitatively, the results are sufficiently similar, and it is unlikely that the outcomes of later analysis interventions (preprocessing, different classifiers, hyperparameter search) were dependent on selecting a particular cross-validation strategy over another.

The selection of classifiers and hyperparameters can lead to an overestimation of the difference in performance between scalars and MoSeq (see Figure 2, Figure 3). Even within the chosen classifiers some preprocessing procedures could remedy the overestimated difference (see Figure 4A-F, H). Similarly, the original hyperparameter search space for the primary linear classifier seemed to not take notice of the limited response when applied to scalar features, as otherwise a larger search space would have been chosen (see Figure 4G).

Finally, an analysis of smaller subsets of the impressively large treatment set shows that the difference between scalar and MoSeq performance, while clearly present and apparently constant, only becomes significant in denser pharmacobehavioral spaces (see Figure 5). Therefore, the advantage of applying MoSeq over scalar measures is more relevant to labs investigating more than 6 to 10 treatment groups in the same study and setup.

In summary, while this publication is rightfully an important contribution to behavioral analysis, accompanied by a large public dataset with a high number of treatment groups, careful analysis reveals a more muted picture of the advantages of MoSeq. Unexpectedly, the choice of measures, parameters, and classifiers led to a significant underestimate of the discriminative performance of basic scalar features. Resolving these issues significantly reduces the gap to the more computationally expensive MoSeq. The topology of scalar and MoSeq features appears at least similar (see Figure 1C, Figure 4F), even if the underlying feature sets have very different properties (see Figure 1A). In effect this could mean that both feature sets are valid transformations of the same underlying, complex behavioral phenotype changes. As locomotion parameters remain easy to explain and reproduce, this raises the question from which point onwards the usage of MoSeq is required for successful analysis of pharmacobehavioral spaces. Considering all of the results presented here, this outcome suggests that the selection of behavioral summary- or classifier models in pre-clinical research should not be solely informed by performance metrics, but also explainability and computational costs, tailored to the specific research questions and treatments used in each study.

## Table of abbreviations

### Abbreviations Full Term

MWU: Mann Whitney U Test
WSR: Wilcoxon Signed-Rank Test

## Code availability statement

The code used to generate the data shown in this study is publicly available. It can be found here: https://github.com/Marti-Ritter/pharmacobehavioral-space-of-moseq-syllables-significantly-overlaps-with-scalar-locomotion-features

## Author Contributions

|  | Conceptualization | Data curation | Formal analysis | Funding acquisition | Investigation | Methodology | Project administration | Resources | Software | Supervision | Validation | Visualization | Writing – original draft | Writing – review & editing |
| --- | --- | --- | --- | --- | --- | --- | --- | --- | --- | --- | --- | --- | --- | --- |
| MR | x | x | x |  | x | x |  |  | x |  | x | x | x | x |
| AB | x |  |  |  |  | x | x |  |  | x |  |  | x | x |

## Generative AI statement

The authors declare that no Generative AI was used in the creation of this manuscript.

